# The complete mitogenome of *Lysmata vittata* (Crustacea: Decapoda: Hippolytidae) and its phylogenetic position in Decapoda

**DOI:** 10.1101/2021.08.04.455109

**Authors:** Longqiang Zhu, Zhihuang Zhu, Leiyu Zhu, Dingquan Wang, Jianxin Wang, Qi Lin

**Author notes:** **(QL)**; **(JW)**; **(ZZ)**.

## Abstract

In this study, the complete mitogenome of *Lysmata vittata* (Crustacea: Decapoda: Hippolytidae) has been determined. The genome sequence was 22003 base pairs (bp) and it included thirteen protein-coding genes (PCGs), twenty-two transfer RNA genes (tRNAs), two ribosomal RNA genes (rRNAs) and three putative control regions (CRs). The nucleotide composition of AT was 71.50%, with a slightly negative AT skewness (−0.04). Usually the standard start codon of the PCGs was ATN, while *cox1*, *nad4L* and *cox3* began with TTG, TTG and GTG. The canonical termination codon was TAA, while *nad5* and *nad4* ended with incomplete stop codon T, and *cox1* ended with TAG. We compared the order of genes of Decapoda ancestor and found that the positions of the two tRNAs genes (*trnA* and *trnR*) of the *L. vittata* were translocated. The phylogenetic tree showed that *L. vittata* was an independent clade, namely Hippolytidae.

## Introduction

*Lysmata vittata* (Crustacea: Decapoda: Hippolytidae) belongs to a small marine ornamental shrimp, commonly known as peppermint shrimp, which is popular in the marine aquarium trade. The species has a special sexual system, ie, protandric simultaneous hermaphrodite (PSH) [1]. It is a member of the clean shrimp family, a common marine ornamental species that originated in the Indian Ocean-Pacific region, including coastal areas such as China, Japan, Philippines and Australia [2–4]. *L. vittata* prefers to move in the range of 2~50 m below the sea surface, usually hiding in the reef during the day and activating at night [5]. In view of the research needs of *L. vittata*, we sequenced its mitogenome sequence.

The mitogenome is a significant tool for studying identification and phylogenetic relationships in the different species [6]. In shrimps, the mitochondria is maternally inherited, usually is circular and approximately 15 to 20 kb in length, including thirteen PCGs, two rRNAs, twenty-two tRNAs and one CR. The mitogenome is a complete system, which not only contains abundant information, but also the phylogenetic tree based on the genome has the advantages of stable and reliable structure.

Decapoda includes the largest number of species in crustaceans (8000 ~ 10000 species), with the greatest economic value and the most widely known invertebrates [7]. It includes many aquatic products with important economic value, such as lobsters, prawns and crabs. Therefore, the phylogeny and classification of decapod crustaceans have been the focus of research for many years. The classification of Hippolytidae was the most controversial family in Decapoda, especially the monophyly of Hippolytidae and the position of the genus *Lysmata* [1, 8]. The Hippolytidae is an important group of marine benthic organisms and a common group in shallow sea biomes. Most species of the Hippolytidae are small shrimps living in shallow water, which are distributed worldwide. It occupies an important position in the animal classification system. However, we are the first to publish the mitochondrial genome sequence of the Hippolytidae species in the GenBank database, which is of great significance for us to expand the database of Hippolytidae.

In this study, the mitogenome of *L. vittata* has been successfully determined, which helps us to understand the characteristics of mtDNA of *L. vittata*. Furthermore, phylogenetic analysis using the nucleotide and amino acid sequences of thirteen PCGs helps us to reconstruct the phylogenetic relationship between *L. vittata* and related species. The addition of newly determined mitogenome complements the record of the mitochondrial gene library of Hippolytidae from scratch.

## Materials and methods

### Mitochondria DNA sequencing and genome assembly

Specimens of *L. vittata* were collected in Xiamen, Fujian province, China. The morphological characteristics of the species follow the previous description of Abdelsalam [9]. Approximately 5g of fresh leaves was harvested for mtDNA isolation using an improved extraction method [10]. After DNA isolation, the isolated DNA was purified according to manufacturer’s instructions (Illumina), and then 1 μg was taken to create short-insert libraries, whose insertion size was 430 bp, followed by sequencing on the Illumina Hiseq 4000 [11] (Shanghai BIOZERON Co., Ltd). The high molecular weight DNA was purified and used for PacBio library prep, BluePippin size selection, then sequenced on the Sequel Squencer.

The raw data obtained by sequencing was processed and then the duplicated sequences were assembled. The mitogenome was reconstructed using a combination of the PacBio Sequel and the Illumina Hiseq data. Assemble the genome framework by the both Illumina and PacBio using SOAPdenovo2.04 [12]. Verifying the assembly and completing the circle or linear characteristic of the mitogenome, filling gaps if there were. Finally, the clean data were mapped to the assembled draft mitogenome to correct the wrong bases, and the most of the gaps were filled through local assembly.

### Validation of mitogenome data

In order to ensure the accuracy of the *L. vittata* mitogenome data, we resequenced the samples on the Illumina HiSeq X10 platform (Nanjing Genepioneer Biotechnologies Co. Ltd).

### Genome annotation and sequence analysis

Mitogenome sequences were annotated using homology-based prediction and de novo prediction, and the EVidenceModeler v1.1 [13] was used to integrate the complete genetic structure. Twenty-two tRNAs and two rRNAs were predicted by tRNAscan-SE [14] and rRNAmmer 1.2 [15]. The circular of the complete *L. vittata* mitogenome graphical map was drawn using OrganellarGenomeDRAW v1.2 [16]. The RSCU of thirteen PCGs (remove incomplete codons) was calculated using MEGA 5.0 [17]. The composition skewness of each component of the genome was calculated according to the following formulas: AT-skew = (A-T) / (A+T); GC-skew = (G-C) / (G+C) [18]. The secondary cloverleaf structure of tRNAs was examined with MITOS WebServer (http://mitos2.bioinf.uni-leipzig.de/index.py) [19].

### Phylogenetic analysis

To reconstruct the phylogenetic relationship among shrimp, the PCGs sequences of the 51 Decapoda species were downloaded from GenBank database (S1 Table). The PCGs sequences of *Euphausia superba* (NC_040987.1) were used as outgroup. The nucleotide and amino acid sequences of 13 PCGs were aligned using MEGA 5.0 [17].

Gblocks was used to identify and selected the conserved regions [20]. Subsequently, Bayesian inference (BI) and Maximum likelihood (ML) analysis were utilized for reconstructing phylogenetic tree by MrBayes v3.2.6 [21] and PhyML 3.1 [22]. According to the Akaike Information Criterion (AIC) [23], GTR + I + G model was considered as the best-fit model for analysis with nucleotide alignments using jModeltest [24], and MtArt + I + G + F model was the optimal model for the amino acid sequence dataset using ProtTest 3.4.2 [25]. In BI analysis, two simultaneous runs of 10000000 generations were conducted for the matrix. Sampling trees every 1000 generations, and diagnostics were calculated every 5000 generations, with three heated and one cold chains to encourage swapping among the Markov-chain Monte Carlo (MCMC) chains. Additionally, the standard deviation of split frequencies was below 0.01 after 10000000 generations, and the potential scale reduction factor (PSRF) was close to 1.0 for all parameters. Posterior probabilities over 0.9 or bootstrap percentage over 75%, the results were regarded as credible [26, 27]. The resulting phylogenetic trees were visualized in Fig Tree v1.4.0.

## Results and discussion

### Genome structure, organization and composition

The mitogenome of *L. vittata* was a typical circular molecule of 22003 bp in size. It contained 37 mitochondrial genes (thirteen PCGs, twenty-two tRNAs, two rRNAs and three CRs) (Fig 1 and S2 Table). Among the 37 genes, the coding direction of the twenty-three genes was clockwise (F-strand), and the coding direction of the remaining fourteen genes was counterclockwise (R-strand) (Fig 1 and S2 Table). The nucleotide composition of the mitogenome was biased toward A and T (T=37.15%, A=34.35%, C=16.69%, G=11.80%) (Table 1). The relatively AT contents of the complete mitogenome were calculated [mitogenome (71.50%), PCGs (69.79%), tRNAs (69.58%) and rRNAs (69.29%)] (Table 1). The AT-skew values (−0.04) and GC-skew values (−0.17) for the entire mitogenome were negative, showing that there were higher Ts than As and Cs than Gs (Table 1). All original sequence data in this study were submitted to the NCBI database under accession number MT478132.

**Fig 1.**
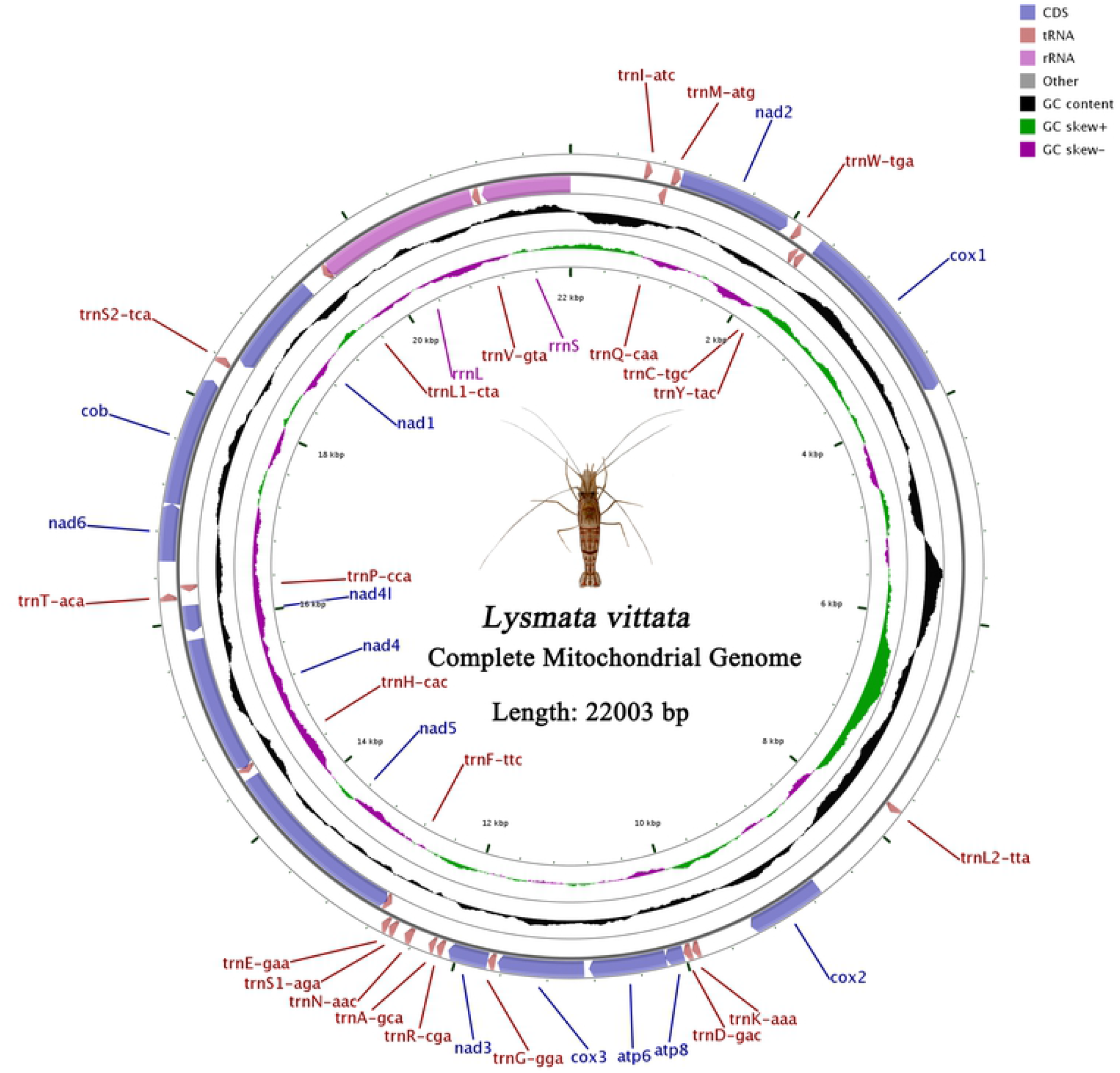
Mitogenome map of *Lysmata vittata*. The genes outside the map were coded on the F strand, whereas the genes on the inside of the map are coded on the R strand. The middle black circle displays the GC content and the inside purple and green circle displays the GC skew.

**Table 1.** Composition and skewness of *Lysmata vittata* mitogenome.

| *Lysmata vittata* | Size(bp) | T (%) | C (%) | A (%) | G (%) | A+T (%) | AT-skew | GC-skew |
| --- | --- | --- | --- | --- | --- | --- | --- | --- |
| Mitogenome | 22003 | 37.15 | 16.69 | 34.35 | 11.80 | 71.50 | -0.04 | -0.17 |
| PCGs | 11144 | 41.09 | 15.25 | 28.70 | 14.96 | 69.79 | -0.18 | -0.01 |
| *atp6* | 675 | 40.15 | 19.41 | 28.30 | 12.15 | 68.44 | -0.17 | -0.23 |
| *atp8* | 165 | 43.64 | 15.76 | 35.15 | 5.45 | 78.79 | -0.11 | -0.49 |
| *cob* | 1137 | 39.40 | 20.14 | 27.88 | 12.58 | 67.28 | -0.17 | -0.23 |
| *cox1* | 1614 | 37.73 | 17.91 | 27.76 | 16.60 | 65.49 | -0.15 | -0.04 |
| *cox2* | 693 | 37.95 | 19.77 | 28.43 | 13.85 | 66.38 | -0.14 | -0.18 |
| *cox3* | 756 | 39.29 | 18.25 | 27.91 | 14.55 | 67.20 | -0.17 | -0.11 |
| *nad1* | 927 | 44.01 | 10.79 | 27.29 | 17.91 | 71.31 | -0.23 | 0.25 |
| *nad2* | 1005 | 43.28 | 18.01 | 29.05 | 9.65 | 72.34 | -0.20 | -0.30 |
| *nad3* | 354 | 42.66 | 18.93 | 26.27 | 12.15 | 68.93 | -0.24 | -0.22 |
| *nad4* | 1336 | 43.11 | 9.51 | 28.59 | 18.79 | 71.70 | -0.20 | 0.33 |
| *nad4l* | 246 | 45.12 | 7.72 | 26.02 | 21.14 | 71.14 | -0.27 | 0.46 |
| *nad5* | 1732 | 41.17 | 9.82 | 31.64 | 17.38 | 72.81 | -0.13 | 0.26 |
| *nad6* | 504 | 44.64 | 17.06 | 28.57 | 9.72 | 73.21 | -0.22 | -0.27 |
| tRNAs | 1512 | 33.27 | 14.02 | 36.31 | 16.40 | 69.58 | 0.04 | 0.08 |
| rRNAs | 2315 | 32.40 | 11.88 | 36.89 | 18.83 | 69.29 | 0.06 | 0.23 |
| CR1 | 650 | 42.15 | 9.85 | 38.31 | 9.69 | 80.46 | -0.05 | -0.01 |
| CR2 | 3821 | 38.50 | 14.39 | 33.73 | 13.37 | 72.23 | -0.07 | -0.04 |
| CR3 | 888 | 42.34 | 13.51 | 34.91 | 9.23 | 77.25 | -0.10 | -0.19 |

### PCGs and codon usage

The PCGs region was 11144 bp long, and accounted 50.6% of the *L. vittata* mitogenome. Nine of thirteen PCGs (*atp6*, *atp8*, *cob*, *cox1-3*, *nad2-3* and *nad6*) were encoded on the light (F) strand, while the other four genes (*nad1*, *nad4L* and *nad4-5*) were encoded on the heavy (R) strand (Table 1). Each PCG was initiated by a canonical ATN codon (ATG for *atp6*, *atp8*, *nad2-5* and *cob*; ATT for *cox2 and nad1*; ATC for *nad6*), except for *cox1* (TTG), *nad4L* (TTG) and *cox3* (GTG) (S2 Table). Two of the thirteen PCGs (*nad5* and *nad4*) terminated with incomplete stop codon T, one PCG (*cox1*) terminated with stop codon TAG, and the other ten PCGs terminated with the canonical termination codon TAA (S2 Table).

The RSCU values of *L. vittata* mitogenome were analyzed and the results were shown in Table 2. The total number of codons in thirteen PCGs was 3714 except eleven canonical stop codons and two incomplete stop codons and the most common amino acids were Ile (AUR) (499), Phe (UUR) (357) and Leu2 (UUR) (315), whereas codons encoding Cys (UGR) (41) and Met (AUR) (24) were rare (Fig 2). The overall A + T content of thirteen PCGs was 69.79%, the AT-skews and GC-skews were negative which implied a higher occurrence of Ts and Cs than As and Gs (Table 1).

**Fig 2.**
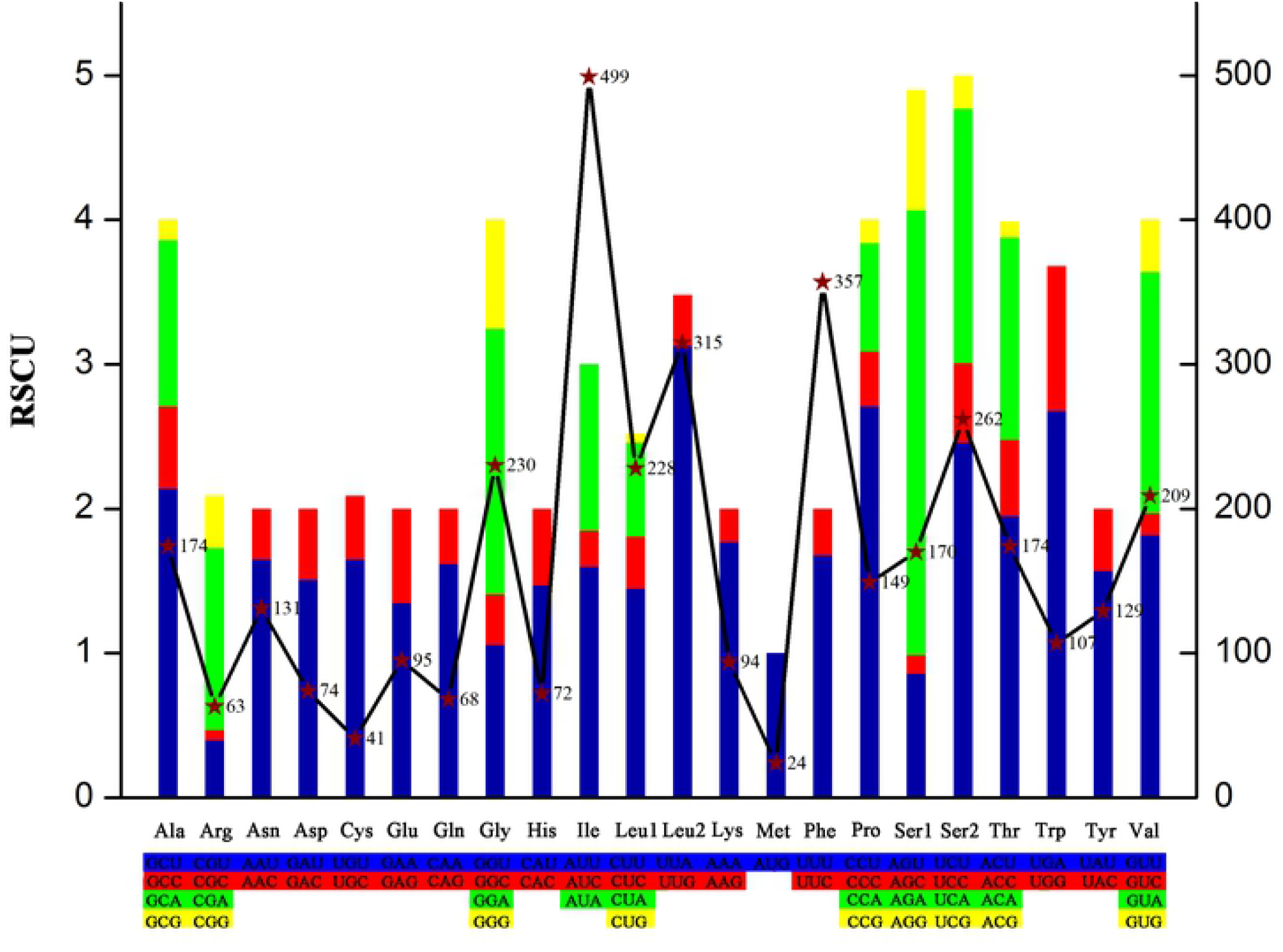
RSCU and Codon distribution in the mitogenome of *L. vittata*. The left ordinate represents RSCU, and the right ordinate represents the number of the Codon distribution.

**Table 2.** The codon number and relative synonymous codon usage (RSCU) in *L. vittata* mitochondrial protein coding genes.

| Codon | Count | RSCU | Codon | Count | RSCU | Codon | Count | RSCU | Codon | Count | RSCU |
| --- | --- | --- | --- | --- | --- | --- | --- | --- | --- | --- | --- |
| UUU(F) | 300 | 1.68 | UCU(S) | 129 | 2.46 | UAU(Y) | 101 | 1.57 | UGU(C) | 32 | 1.56 |
| UUC(F) | 57 | 0.32 | UCC(S) | 29 | 0.55 | UAC(Y) | 28 | 0.43 | UGC(C) | 9 | 0.44 |
| UUA(L) | 283 | 3.13 | UCA(S) | 92 | 1.76 | UAA(*) | 10 | 0.29 | UGA(W) | 92 | 2.68 |
| UUG(L) | 32 | 0.35 | UCG(S) | 12 | 0.23 | UAG(*) | 1 | 0.03 | UGG(W) | 15 | 1 |
| CUU(L) | 131 | 1.45 | CCU(P) | 101 | 2.71 | CAU(H) | 53 | 1.47 | CGU(R) | 12 | 0.4 |
| CUC(L) | 33 | 0.36 | CCC(P) | 14 | 0.38 | CAC(H) | 19 | 0.53 | CGC(R) | 2 | 0.07 |
| CUA(L) | 59 | 0.65 | CCA(P) | 28 | 0.75 | CAA(Q) | 55 | 1.62 | CGA(R) | 38 | 1.26 |
| CUG(L) | 5 | 0.06 | CCG(P) | 6 | 0.16 | CAG(Q) | 13 | 0.38 | CGG(R) | 11 | 0.36 |
| AUU(I) | 266 | 1.6 | ACU(T) | 85 | 1.95 | AAU(N) | 108 | 1.65 | AGU(S) | 45 | 0.86 |
| AUC(I) | 42 | 0.25 | ACC(T) | 23 | 0.53 | AAC(N) | 23 | 0.35 | AGC(S) | 7 | 0.13 |
| AUA(I) | 191 | 1.15 | ACA(T) | 61 | 1.40 | AAA(K) | 83 | 1.77 | AGA(S) | 93 | 3.08 |
| AUG(M) | 24 | 1 | ACG(T) | 5 | 0.11 | AAG(K) | 11 | 0.23 | AGG(S) | 25 | 0.83 |
| GUU(V) | 95 | 1.82 | GCU(A) | 93 | 2.14 | GAU(D) | 56 | 1.51 | GGU(G) | 61 | 1.06 |
| GUC(V) | 8 | 0.15 | GCC(A) | 25 | 0.57 | GAC(D) | 18 | 0.49 | GGC(G) | 20 | 0.35 |
| GUA(V) | 87 | 1.67 | GCA(A) | 50 | 1.15 | GAA(E) | 64 | 1.35 | GGA(G) | 106 | 1.84 |
| GUG(V) | 19 | 0.36 | GCG(A) | 6 | 0.14 | GAG(E) | 31 | 0.65 | GGG(G) | 43 | 0.75 |

### Transfer RNAs and Ribosomal RNAs

The mitogenome of *L. vittata* contained twenty-two tRNAs and these genes ranged from 60 (*trnA*) to 77 bp (*trnN*) (S2 Table). The tRNAs showed a strong A +T bias (69.58%), while they also exhibited positive AT-skew (0.04) and GC-skew (0.08) (Table 1). Eight tRNAs [*trnQ* (CAA), *trnC* (UGC), *trnY* (UAC), *trnF* (UUC), *trnH* (CAC), *trnP* (CCA), *trnL1* (CUA) and *trnV* (GUA)] were present on the R strand and the remaining fourteen were present on the F strand (S2 Table). The examined secondary structure of twenty-two tRNAs was shown in S1 Fig. The other twenty-one tRNAs had typical cloverleaf secondary structure except that *trnS1* (AGA) lacked the dihydropyridine (DHU) arm [18, 19, 27, 28] (S1 Fig). In the secondary structure of the tRNAs, the most common non-Watson–Crick base pair was G–U (e.g. *trnC*, *trnE*), followed by U–U (e.g. *trnA*, *trnC*) [19]. In addition, several mismatches were common in tRNAs, such as A–C (e.g. *trnA*), C–U (e.g. *trnA*, *trnG*) and A–A (e.g. *trnM*, *trnS1*) (S1 Fig).

Two rRNA genes were found on the R strand. The *rrnL* was 1494 bp and *rrnS* was 821 bp, one located between *trnL1* and *trnV* and another located between *trnV* and CR1 (S2 Table and Fig 1). The total A+T content of the two rRNAs was 69.29%, with a positive AT-skew (0.06) (Table 1).

### Overlapping and intergenic regions

The mitogenome of *L. vittata* contained four overlapping regions, these four pairs of genes were presented: *atp8* / *atp6*, *trnE* / *trnF*, *nad4* / *nad4L* and *trnL1* / *rrnL*, with the longest 23 bp overlap located between *trnL1* and *rrnL* (S2 Table). The 27 intergenic regions were found with a length varying from 2 ~ 3821 bp (S2 Table). Three putative CRs had been identified in *L. vittata* mitogenome. The CR1 was located between *rrnS* and *trnI*, with a length of 650 bp, and the A+T content was 80.46%. The CR2 was located between *cox1* and *trnL2*, with a length of 3821 bp, and the A+T content was 72.23%. The CR3 was located between *trnL2* and *cox2*, with a length of 888 bp, and the A+T content was 77.25% (Table 1 and S2 Table).

To our knowledge, this study is the first reported mitogenome from the genus *Lysmata*. How multiple CRs were generated and evolved in the mitogenome of *Lysmata* is a novel problem that has not yet been solved, and more mitogenomes of *Lysmata* are still needed to clarify the mechanism forming this phenomenon.

### Gene rearrangement

Compared with the gene order of a Decapoda ancestor [20, 29], two tRNA gene (*trnA* and *trnR*) positions of *L. vittata* had translocated, which indicates that the *L. vittata* was quite unconserved in its evolution (Fig 3). In fact, gene rearrangement was a very common phenomenon in the mitogenome and the rearrangement mainly occurred in tRNA genes. Gene arrangement was stable, and it could be used as an important phylogenetic marker in the analysis of evolutionary perspective on shrimp. At present, no other species in the Hippolytidae have been tested for mitogenome, and the common characteristics of gene order were not easy to determine.

**Fig 3.**
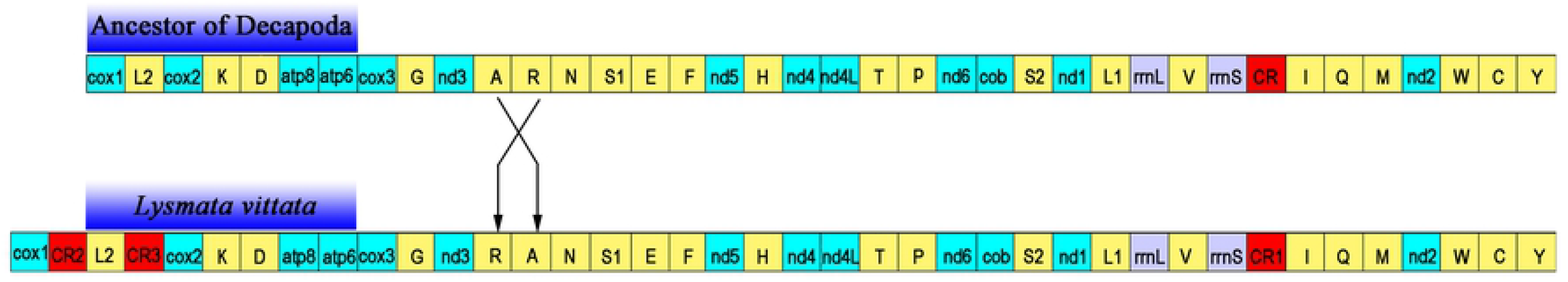
Comparison of the order of mitochondrial genes of *Lysmata vittata* and the ancestor of Decapoda.

### Phylogenetic analysis

Using ML and BI analysis methods, phylogenetic analysis was performed based on the nucleotide and amino acid sequences of thirteen PCGs of the species in S1 Table, and the analysis results were presented (Fig 4 and Fig 5). The phylogenetic tree based on the nucleotide sequence of thirteen PCGs showed that the monophyly of each family was basically well supported, especially the clade of the Hippolytidae was strongly supported (ML BP = 100%; BI PP = 1). A basal split separates two clades, with insignificant support (Fig 4). The first clade revealed the two phylogenetic relationships: (Hippolytidae + (Atyidae + (Alpheidae + Palaemonidae))) and (Palinuridae + (Astacidae + (Nephropsidae + Enoplometopidae))). The second clade revealed the one phylogenetic relationship: (Sergestidae + (Solenoceridae + Penaeidae)) (Fig 4). The phylogenetic tree based on the amino acid sequence of 13 PCGs revealed that the phylogenetic relationship between Hippolytidae and Atyidae has changed as follows: (Atyidae + (Hippolytidae + (Alpheidae + Palaemonidae))). However, the clade of the Hippolytidae was very weak support (ML BP = 52%; BI PP = 0) (Fig 5). We could still reach a conclusion that the Hippolytidae was an older family than Atyidae, and the Atyidae formed a sister group to Alpheidae – Palaemonidae. The Caridea were dominated by Palaemonidae, followed by Alpheidae, Atyidae and Hippolytidae [30]. At present, the phylogenetic study of the Hippolytidae was limited to the partial fragments of mitochondrial genes *16S* or *12S* of individual species in several genera (such as *Lysmata*, *Exhippolysmata*, *Ligur*, *Mimocaris* and *Lysmatella*) [31–34]. The successful determination of the mitogenome of *L. vittata* could provide a deeper understanding of the phylogenetic status of the Hippolytidae.

**Fig 4.**
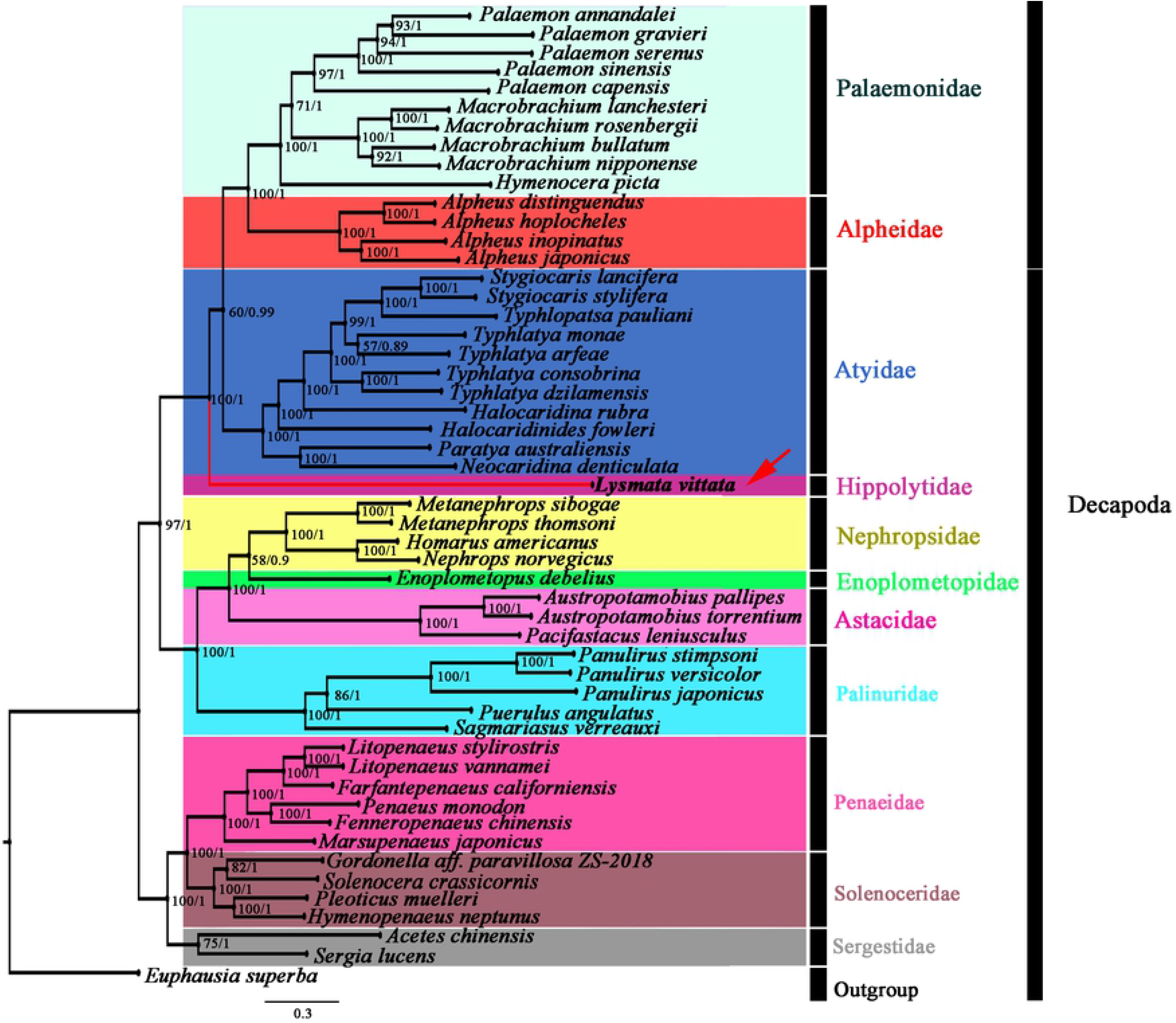
Phylogenetic tree inferred from nucleotide sequences of 13 PCGs of the mitogenome using ML and BI methods (BP / PP).

**Fig 5.**
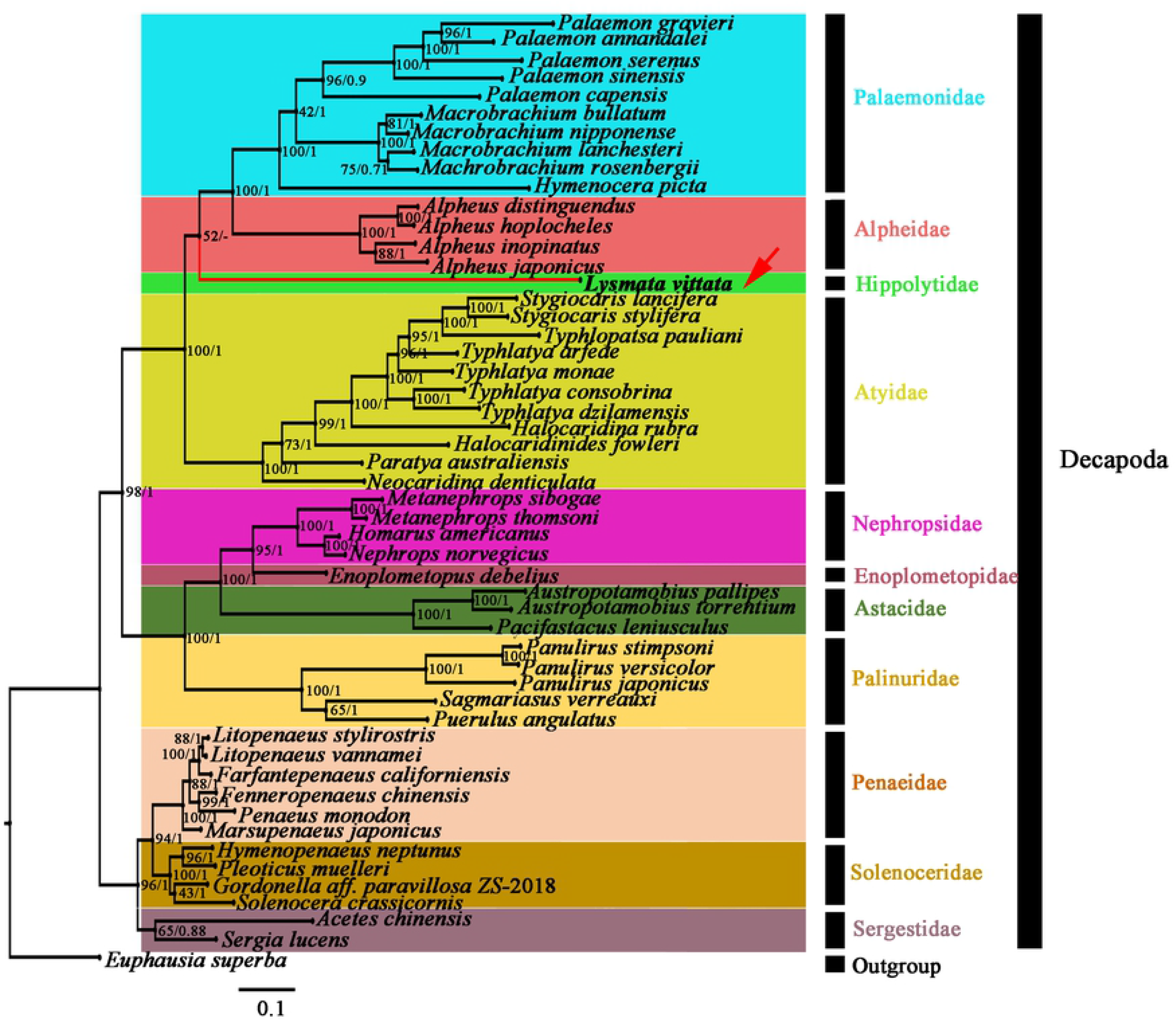
Phylogenetic tree inferred from amino acid sequences of 13 PCGs of the mitogenome using ML and BI methods (BP / PP).

## Conclusion

In this study, we successfully obtained the mitogenome sequence of the *L. vittata*, which was also the first species of the Hippolytidae to publish the mitogenome sequence in the GenBank database. The genome sequence was 22003 base pairs (bp) and it included 37 genes and three CRs. Each PCGs was initiated by a canonical ATN codon, except for *cox1*, *nad4L* and *cox3*, which were initiated by a TTG, TTG and GTG. Two of the thirteen PCGs (*nad5* and *nad4*) terminated with incomplete stop codon T, and one (*cox1*) terminated with stop codon TAG. The AT-skew (−0.04) and the GC-skew (−0.17) were both negative in the mitogenomes of *L. vittata*. Compared with the gene order of a Decapoda ancestor, the gene arrangement order of the *L. vittata* has changed. Futhermore, phylogenetic analyses showed that *L. vittata* was not in the clades of other families, but was an independent clade, namely the Hippolytidae.

## Supporting information

**S1 Table. List of species used to construct the phylogenetic tree.**

(DOC)

**S2 Table. Summary of *Lysmata vittata* mitogenome.**

(DOC)

**S1 Fig. Predicted secondary structure for the tRNAs of Lysmata vittata mitogenome.**

(TIF)

## Acknowledgements

The authors thank the parents for their material and spiritual support.

## Author contributions

**Supervision:** Zhihuang Zhu, Qi Lin, Jianxin Wang.

**Funding acquisition:** Zhihuang Zhu, Qi Lin, Jianxin Wang.

**Methodology:** Longqiang Zhu, Leiyu Zhu, Dingquan Wang.

**Software:** Longqiang Zhu.

**Writing - original draft:** Longqiang Zhu.

**Writing - review & editing:** Zhihuang Zhu, Qi Lin, Jianxin Wang.

## Data Availability Statement

Data are available from the NCBI database (accession number MT478132).

## References

1. De Grave S, Li C P, Tsang L M, Chan T Y. Unweaving hippolytoid systematics (Crustacea, Decapoda, Hippolytidae): resurrection of several families. Zoologica Scripta. 2014; 43: 496–507. https://doi.org/10.1111/zsc.12067

2. Chace F A, Jr. The Caridean Shrimps (Crustacea: Decapoda) of the Albatross Philippine Expedition, 1907-1910, Part 7: Families Atyidae, Eugonatonotidae, Rhynchocinetidae, Bathypalaemonellidae, Processidae, and Hippolytidae. Smithsonian Contributions to Zoology. 1997; 587: 1–106. https://doi.org/10.5479/si.00810282.381.1

3. Ahyong S T. New species and new records of Caridea (Hippolytidae: Pasiphaeidae) from New Zealand. Zootaxa. 2010; 341–357. https://doi.org/10.1163/156854009X427333

4. Okuno, J. Lysmata lipkei, A new species of peppermint shrimp (Decapoda, Hippolytidae) from warm temperate and subtropical waters of Japan. Studies on Malacostraca. 2010; 597–610. https://doi.org/10.1163/9789047427759_042

5. Marin I N, Korn O M, Kornienko E S. (2012) The caridean shrimp Lysmata vittata (Stimpson, 1860) (Decapoda: Hippolytidae): A new species for the fauna of Russia. Russian Journal of Marine Biology. 2012; 38. https://doi.org/10.1134/S1063074012040062

6. Zheng N, Sun Y X, Yang L L, Wu L, Muhammad AN, Chen C et al. Characterization of the complete mitochondrial genome of Biston marginata (Lepidoptera: Geometridae) and phylogenetic analysis among lepidopteran insects. International Journal of Biological Macromolecules. 2018; S0141-8130(17): 32480–32487. https://doi.org/10.1016/j.ijbiomac.2018.02.110 PMID: 29462677

7. Shan D N. Crustacean Biology, Beijing: Science Press. 1993.

8. De Grave S, Fransen C H J M. Carideorum Catalogus: The recent species of the dendrobranchiate, stenopodidean, procarididean and caridean shrimps. Zoologische Mededelingen. 2011; 85: 195–589.

9. Abdelsalam K M. First record of the exotic lysmatid shrimp Lysmata vittata (Stimpson, 1860) (Decapoda: Caridea: Lysmatidae) from the Egyptian Mediterranean coast. Mediterranean Marine Science. 2018; 19: 124–131. https://doi.org/10.12681/mms.15591

10. Chen J, Guan R, Chang S, Du T, Zhang H, Xing H. Substoichiometrically different mitotypes coexist in mitochondrial genomes of Brassica napus L. PLoS One. 2011; 6: e17662. https://doi.org/10.1371/journal.pone.0017662 PMID: 21423700

11. Borgstrom E, Lundin S, Lundeberg J. Large scale library generation for high throughput sequencing. PLoS One. 2011; 6: e19119. https://doi.org/10.1371/journal.pone.0019119 PMID:21589638

12. Luo R, Liu B, Xie Y, Li Z, Huang W, Yuan J et al. SOAPdenovo2: an empirically improved memory-efficient short-read de novo assembler. Gigascience. 2012; 1(1): 18. https://doi.org/10.1186/2047-217X-1-18 PMID: 23587118

13. Haas B J, Salzberg S L, Zhu W, Pertea M, Allen J E, Orvis J et al. Automated eukaryotic gene structure annotation using EVidenceModeler and the Program to Assemble Spliced Alignments. Genome Biology. 2008; 9(1): R7. https://doi.org/10.1186/gb-2008-9-1-r7 PMID: 18190707

14. Lowe T M, Eddy S R. tRNAscan-SE: a program for improved detection of transfer RNA genes in genomic sequence. Nucleic Acids Res. 1997; 25: 955–964. https://doi.org/10.1093/nar/25.5.955 PMID: 9023104

15. Lagesen K, Hallin P, Rodland E A, Staerfeldt H H, Rognes T, Usser D W. RNAmmer: consistent and rapid annotation of ribosomal RNA genes. Nucleic Acids Res. 2007; 35: 3100–3108. https://doi.org/10.1093/nar/gkm160 PMID: 17452365

16. Lohse M, Drechsel O, Bock R. OrganellarGenomeDRAW (OGDRAW): a tool for the easy generation of high-quality custom graphical maps of plastid and mitochondrial genomes. Current Genetics. 2007; 52: 267–274. https://doi.org/10.1007/s00294-007-0161-y PMID: 17957369

17. Tamura K, Peterson D, Peterson N, Stecher G, Nei M, Kumar S. MEGA5: Molecular Evolutionary Genetics Analysis Using Maximum Likelihood, Evolutionary Distance, and Maximum Parsimony Methods. Molecular Biology & Evolution. 2011; 28: 2731–2739. https://doi.org/10.1093/molbev/msr121 PMID: 21546353

18. Yang Z H, Yang T T, Liu Y, Zhang H B, Tang B P, Liu Q N et al. The complete mitochondrial genome of Sinna extrema (Lepidoptera: Nolidae) and its implications for the phylogenetic relationships of Noctuoidea species. International Journal of Biological Macromolecules. 2019; 137: 317–326. https://doi.org/10.1016/j.ijbiomac.2019.06.238 PMID: 31265851

19. Wang Z, Wang Z, Shi X, Wu Q, Tao Y, Guo H et al. Complete mitochondrial genome of Parasesarma affine (Brachyura: Sesarmidae): Gene rearrangements in Sesarmidae and phylogenetic analysis of the Brachyura. International Journal of Biological Macromolecules. 2018; S0141-8130(18): 32224–4. https://doi.org/10.1016/j.ijbiomac.2018.06.056 PMID: 29908270

20. Castresana J. Selection of conserved blocks from multiple alignments for their use in phylogenetic analysis. Mol Biol Evol. 2000; 17: 540–552. https://doi.org/10.1093/oxfordjournals.molbev.a026334 PMID: 10742046

21. Ronquist F, Huelsenbeck J, Teslenko M. Draft MrBayes version 3.2 Manual: Tutorials and Model Summaries. Scarcelli. 2011; 1-103.

22. Guindon S, Gascuel O. PhyML: “A simple, fast and accurate algorithm to estimate large phylogenies by maximum likelihood”. Systematic Biology. 2003; 52: 696–704.

23. Yamaoka K, Nakagawa T, Uno T. Application of Akaike’s information criterion (AIC) in the evaluation of linear pharmacokinetic equations. Journal of Pharmacokinetics and Biopharmaceutics. 1978; 6: 165–175. https://doi.org/10.1007/BF01117450 PMID: 671222

24. Darriba D, Taboada G L, Doallo R, Posada D. jModelTest 2: more models, new heuristics and high-performance computing. Nature Methods. 2012; 9: 772. https://doi.org/10.1038/nmeth.2109 PMID: 22847109

25. Abascal F, Zardoya R, Posada D. ProtTest: selection of best-fit models of protein evolution. Bioinformatic. 2005; 21: 2104–2105. https://doi.org/10.1093/bioinformatics/bti263 PMID: 15647292

26. Hillis D M, Bull J J. An Empirical Test of Bootstrapping as a Method for Assessing Confidence in Phylogenetic Analysis. Systematic Biology. 1993; 42: 182–192. https://doi.org/10.2307/2992540

27. Zhu X Y, Xin Z Z, Liu Y, Wang Y, Huang Y, Yang Z H et al. The complete mitochondrial genome of Clostera anastomosis (Lepidoptera: Notodontidae) and implication for the phylogenetic relationships of Noctuoidea species. International Journal of Biological Macromolecules. 2018; S0141-8130(18): 32119–32126. https://doi.org/10.1016/j.ijbiomac.2018.06.188 PMID: 29981329

28. Zhu X Y, Xin Z Z, Wang Y, Zhang H B, Zhang D Z, Wang Z F et al. The complete mitochondrial genome of Clostera anachoreta (Lepidoptera: Notodontidae) and phylogenetic implications for Noctuoidea species. Genomics. 2017; S0888-7543(17): 30025–30033. https://doi.org/10.1016/j.ijbiomac.2018.06.188 PMID: 28435087

29. Shen X, Ren J, Cui Z, Sha Z, Wang B, Xiang J et al. The complete mitochondrial genomes of two common shrimps (Litopenaeus vannamei and Fenneropenaeus chinensis) and their phylogenomic considerations. Gene. 2007; 403: 98–109. https://doi.org/10.1016/j.gene.2007.06.021 PMID: 17890021

30. Grave S D, Fransen, C H J M. Carideorum Catalogus: The recent species of the dendrobranchiate, stenopodidean, procarididean and caridean shrimps (Crustacea: Decapoda). Zoologische Mededelingen. 2011; 85: 195–589.

31. Alves D F R, Lima D J M, Hirose G L, Martinez P A, Dolabella S S, Barros-Alves S D P. Morphological and molecular analyses confirm the occurrence of two sympatric Lysmata shrimp (Crustacea, Decapoda) in the southwestern Atlantic. Zootaxa. 2018; 4526: 41–55. https://doi.org/10.11646/zootaxa.4526.1.3 PMID: 30486089

32. Baeza J A, Fuentes M S. Exploring phylogenetic informativeness and nuclear copies of mitochondrial DNA (numts) in three commonly used mitochondrial genes: mitochondrial phylogeny of peppermint, cleaner, and semi-terrestrial shrimps (Caridea: Lysmata, Exhippolysmata, and Merguia). Zoological Journal of the Linnean Society. 2013; 168: 699–722. https://doi.org/10.1111/zoj.12044

33. Baeza J A. Molecular phylogeny of broken-back shrimps (genus Lysmata and allies): A test of the ‘Tomlinson–Ghiselin’ hypothesis explaining the evolution of hermaphroditism. Molecular Phylogenetics and Evolution. 2013; 69: 46–62. https://doi.org/10.1016/j.ympev.2013.05.013 PMID: 23727055

34. Baeza J A. Molecular systematics of peppermint and cleaner shrimps: phylogeny and taxonomy of the genera Lysmata and Exhippolysmata (Crustacea: Caridea: Hippolytidae). Zoological Journal of the Linnean Society. 2010; 160: 254–265. https://doi.org/10.1111/j.1096-3642.2009.00605.x

